# Selected plasma-associated molecular species remodel the ABCB1 linker with effects retained under structure-derived tethering

**DOI:** 10.1101/2025.08.13.670033

**Authors:** Mansoureh Hoseini, Hossein Abbasi

## Abstract

Multidrug resistance mediated by P-glycoprotein/ABCB1 remains a major limitation to chemotherapy, and cold atmospheric plasma has been reported to restore drug sensitivity in resistant cancer models. However, the molecular pathways connecting plasma-associated species to P-glycoprotein regulation remain unclear. Here, we used a two-stage reduced-model strategy to test whether selected plasma-associated molecular species can noncovalently remodel the intrinsically disordered NBD1–TMD2 linker of human ABCB1 and whether phenotypes detected in an unconstrained screening model persist when linker terminal geometry is restricted to a structure-derived range. We first screened the free linker against H_2_O_2_, NO, or O_2_ at three molecular loadings and then repeated the same perturbations under terminal tethering, yielding sixty 100-ns all-atom molecular-dynamics trajectories. The free linker showed species-and loading-dependent chemical susceptibility: high-loading H_2_O_2_ produced a coherent restriction-oriented phenotype, whereas low-loading O_2_ drove a pronounced compact, low-exchange redistribution. Chemical responsiveness was not eliminated by tethering. High-loading H_2_O_2_ retained a compact, low-exchange phenotype, and high-loading O_2_ narrowed the projected conformational ensemble and reduced selected exchange modes. In contrast, the strong low-loading O_2_ response was largely suppressed, indicating dependence on conformational routes enabled by unconstrained terminal motion. Across conditions, responses were spatially heterogeneous; total species–linker attraction did not predict remodeling magnitude, whereas intramolecular H-bond-network reorganization tracked conformational restriction more closely. Together, these simulations establish chemical susceptibility of the ABCB1 linker while distinguishing mechanically persistent from free-motion-dependent responses. The persistent phenotypes provide candidates for testing in full-length transporter simulations and functional assays of ATPase–transport coupling and drug efflux.

## 1. Introduction

Multidrug resistance remains a major cause of chemotherapy failure and continues to limit durable responses across many malignancies. Among the mechanisms contributing to this phenotype, overexpression of P-glycoprotein (P-gp/ABCB1) is particularly important because this transporter reduces the intracellular accumulation of numerous structurally unrelated anticancer agents. P-gp is an ATP-driven efflux pump of the ATP-binding cassette superfamily that couples nucleotide binding and hydrolysis to large-scale conformational rearrangements and substrate export [1–5]. Despite decades of investigation, direct pharmacological inhibition of P-gp has achieved limited clinical success because of toxicity, insufficient selectivity, unfavorable drug–drug interactions, and interference with the transporter’s physiological protective functions [6–10]. These limitations motivate complementary strategies that target the internal dynamics supporting transport rather than simply blocking the substrate-binding cavity.

P-gp consists of two homologous halves, each containing a transmembrane domain and a cytosolic nucleotide-binding domain. Productive transport requires coordinated communication between these domains as the protein progresses through inward-facing, nucleotide-bound, and outward-facing conformations [11–16]. A flexible cytosolic linker connecting nucleotide-binding domain 1 (NBD1) to transmembrane domain 2 (TMD2) occupies a strategically important position within this coupling network. The linker is unresolved in most experimental structures, consistent with an intrinsically disordered and conformationally heterogeneous ensemble. Nevertheless, it is not a passive spacer. Experimental perturbation of this region alters ATP hydrolysis and drug transport, and its structural flexibility is required for efficient P-gp function [17,18]. Molecular-dynamics studies have further implicated the linker in stabilizing nucleotide-binding-domain arrangements and modulating communication between the two halves of the transporter [19–22].

Unlike a folded regulatory domain, an intrinsically disordered linker functions through a population of interconverting conformations rather than a single dominant structure. A molecular perturbation may therefore be expressed as redistribution of state populations, altered exchange kinetics, or localized changes in flexibility without formation of a new folded state [23–25]. This ensemble-based behavior makes the ABCB1 linker a plausible target for weak and transient interactions that can bias conformational populations without requiring a stable binding complex.

This possibility is relevant to cold atmospheric plasma (CAP), an emerging anticancer modality that produces reactive oxygen and nitrogen species, charged particles, electric fields, and other physicochemical stimuli [26–29]. CAP and plasma-activated media have shown selective antitumor activity and have enhanced chemotherapy responses in several resistant cancer-cell models [30–34]. Proposed mechanisms include oxidative stress, membrane damage, altered gene expression, redox signaling, and activation of cell-death pathways. Whether CAP-associated molecular species can also directly perturb the conformational dynamics of protein regions involved in drug transport remains largely unexplored.

The NBD1–TMD2 linker therefore provides a plausible molecular entry point for CAP-associated noncovalent perturbation. In an intrinsically disordered ensemble, conformational consequences need not scale with average interaction strength: transient contacts can preferentially stabilize or destabilize pre-existing states, while stronger but more distributed interactions may produce little coherent remodeling. Linker remodeling would not by itself demonstrate altered ATPase activity, drug efflux, or reversal of multidrug resistance. It would instead identify a specific molecular step through which CAP-associated chemistry could plausibly influence the interdomain coordination required for P-gp transport.

Hydrogen peroxide, nitric oxide, and molecular oxygen were selected as a chemically distinct test set. H_2_O_2_ is polar and capable of electrostatic and hydrogen-bonding interactions, whereas NO and O_2_ are substantially less polar and are expected to interact predominantly through dispersion forces. These molecules represent chemically distinct, relatively long-lived components associated with CAP exposure [35]. Comparing them at several finite-copy-number loadings therefore permits assessment of whether linker remodeling depends on molecular identity and loading rather than on a single generic measure of interaction strength.

A response observed in an isolated peptide fragment is not automatically transferable to the linker embedded within ABCB1. An unconstrained chain can collapse, extend, or fold back through geometries that may become less accessible when its termini are connected to neighboring domains; terminal constraints can likewise reshape accessible configurations without uniformly immobilizing the intervening sequence [23–25]. We therefore treated terminal mechanics as a transferability test. The free-linker model (FL) provided a permissive screen for chemical susceptibility, whereas the tethered-linker model (TL) imposed a structure-derived terminal-separation range while preserving internal fluctuations. The two models were not intended to represent alternative physiological states; their comparison was used to distinguish chemical phenotypes that depend strongly on unrestricted terminal motion from those retained or emerging under a mechanically constrained reduced representation.

Accordingly, we used all-atom molecular dynamics to ask two sequential questions: whether selected CAP-associated molecular species can remodel the human ABCB1 NBD1–TMD2 linker, and whether those chemical phenotypes remain detectable after imposing structure-derived terminal tethering. FL was screened independently with H_2_O_2_, NO, or O_2_ at three molecular loadings, after which the same perturbations were evaluated in TL. The resulting ensembles were characterized using global conformational descriptors, state-matched treatment effects, Bayesian difference-in-differences interactions, projected radius-of-gyration–end-to-end-distance landscapes, residue-resolved RMSF and backbone order, intramolecular hydrogen bonding, and comparative species–linker interaction energies. We hypothesized that molecular identity and loading would produce distinct changes in linker populations and exchange dynamics, and that mechanically persistent responses would identify higher-priority candidates for testing in full-length ABCB1.

## 2. Materials and Methods

### 2.1. Linker model and two-stage simulation design

The study focused on the intrinsically disordered linker connecting nucleotide-binding domain 1 (NBD1) to transmembrane domain 2 (TMD2) in human P-glycoprotein/ABCB1 (Figure 1A). Because this segment is unresolved experimentally, residues 628–711 were extracted from the AlphaFold2 model of human ABCB1 (AlphaFold DB accession AF-P08183-F1; UniProt P08183) and used as the initial linker conformation [36]. The design followed a sequential reduced-model strategy: FL served as the permissive screening model, and the same species and loadings were subsequently evaluated in TL to test whether detected chemical responses persisted when terminal separation was restricted to a structure-derived range. FL and TL were not treated as alternative biological states of ABCB1.

**Figure 1.**
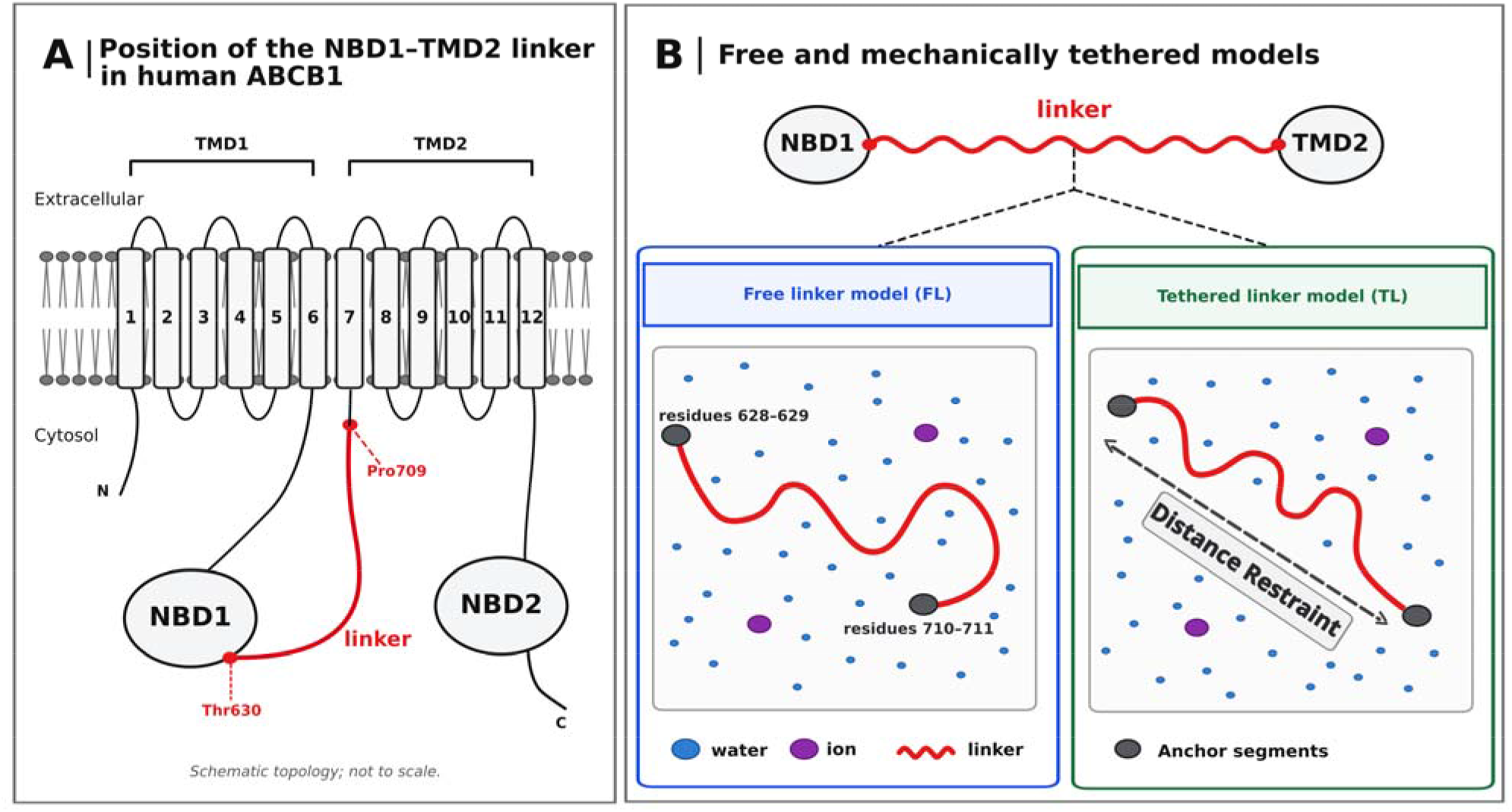
Two-stage reduced-model strategy for testing CAP-associated remodeling of the ABCB1 linker. **(A)**, Schematic position of the Thr630–Pro709 NBD1–TMD2 linker in human ABCB1. **(B)**, FL and TL models. Both FL and TL contain residues 628–711; only residues 630–709 (the intervening red chain) were included in linker analyses. FL serves as the permissive screening model in which terminal separation is unconstrained. TL introduces four pairwise Cα distance restraints between residues 628–629 and 710–711 to test whether chemical responses persist when terminal motion is restricted to a structure-derived range. The permitted restraint interval is 7.0–8.0 nm, with restoring potentials outside this interval.

Both the FL and TL systems contained the identical ABCB1 segment spanning residues 628–711 (Figure 1B). Residues Thr630–Pro709 were defined a priori as the linker analysis region, whereas residues 628–629 and 710–711 were retained as flanking anchor segments and excluded from linker-level conformational and residue-resolved analyses. The two models differed only in their terminal boundary condition. In the free-linker model, no restraint was applied to the anchor segments. TL contained four pairwise type-10 distance restraints connecting the Cα atoms of residues 628–629 to those of residues 710–711. Each restraint had a zero-force interval of 7.0–8.0 nm, a harmonic region from 8.0 to 8.5 nm and below 7.0 nm, and a linear regime above 8.5 nm, with k_dr_ = 800 kJ mol^-1^ nm^-2^. This structure-derived pairwise terminal restraint approximated the attachment to neighboring ABCB1 domains while retaining internal linker fluctuations. The permitted range was selected to encompass anchor separations observed in available P-glycoprotein structures^[11–13]^.

All simulations were performed with GROMACS 2024.2 ^[37]^ and the CHARMM36m force field ^[38]^. Each construct was solvated in a cubic TIP3P water box of approximately 1945 nm^3^ (about 12.5 nm per side and 62,950 water molecules), neutralized with counterions, and simulated under periodic boundary conditions. Hydrogen peroxide (H_2_O_2_), nitric oxide (NO), and molecular oxygen (O_2_) were represented using validated force-field parameters ^[35,39–42]^.

Each species was examined individually at loadings of 1, 5, and 10 molecules. These systems corresponded to nominal concentrations of approximately 0.85, 4.3, and 8.5 mM, respectively, based on the mean simulation-box volume. These finite-copy-number loadings are operational simulation conditions and are not intended as direct equivalents of cellular exposure concentrations. Molecules were placed randomly in the solvent while avoiding steric overlap. A species-free control was generated for each mechanical state. The design therefore comprised 20 conditions—two controls and 18 single-species treatments—with three independent 100-ns replicas per condition, yielding 60 trajectories and 6.0 μs of aggregate production sampling.

### 2.2. Molecular dynamics protocol

Systems were energy-minimized by steepest descent, equilibrated for 1 ns in the NVT ensemble at 310.15 K using the Bussi–Donadio–Parrinello stochastic velocity-rescaling thermostat ^[43]^, and then equilibrated for 1 ns in the NPT ensemble at 1 atm using stochastic cell rescaling ^[44]^. Long-range electrostatics were treated with particle-mesh Ewald summation, and a 1.2-nm real-space cutoff was applied to nonbonded interactions. Bonds involving hydrogen were constrained with LINCS, permitting a 2-fs integration step.

The first 50 ns of each production trajectory were excluded, and quantitative analyses used the final 50 ns. Replica-level summaries—not individual frames—were treated as the independent observations in condition-level inference.

### 2.3. Global conformational and mechanical descriptors

Seven complementary descriptors were used to characterize global linker behavior: conformational entropy, covariance trace, Cα root-mean-square fluctuation (RMSF), effective stiffness, the minimum backbone order parameter (S^2^_min_), persistence length, and backbone dihedral-transition count.

Conformational entropy was estimated from the mass-weighted covariance matrix using the Schlitter approximation ^[45]^ after least-squares removal of translation and rotation. The covariance trace, tr(C), quantified the total amplitude of collective positional fluctuations. Residue-level Cα RMSF was calculated after structural alignment, and the mean across residues 630–709 was used as the global RMSF measure. Backbone orientational dynamics were summarized by the minimum residue-level generalized order parameter, *S*^2^_min_; larger values indicate greater orientational order. Persistence length was obtained from the decay of directional correlations along the backbone.

The end-to-end distance, r, was calculated between the terminal groups used to define the TL restraint. For FL, the one-dimensional potential of mean force was calculated from the sampled distance distribution,

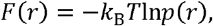

and the region around the minimum was fit to

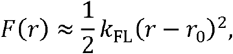

to estimate the effective spring constant *k*_FL_. For TL, the flat-bottom window precluded an equivalent PMF fit. Effective stiffness, *k*_TL_, was therefore estimated from end-to-end fluctuations using the equipartition theorem:

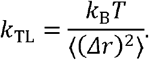

Because the two estimates arise from different mechanical reference models, stiffness was used only for treatment-versus-control comparisons within FL or TL and was excluded from formal cross-boundary interaction analysis.

Backbone dihedral transitions were quantified with gmx chi using the default GROMACS rotamer-core definition. Transitions were counted from changes in rotamer assignment in equally spaced frames without additional smoothing or a minimum residence-time requirement. The total number of transitions over the analyzed 50-ns window was used as a measure of local conformational exchange.

### 2.4. State-matched effects and boundary-condition interactions

Each treatment was compared with its state-matched control using small-sample-corrected Hedges *g*,

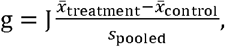

where *J* is the bias correction and s_pooled_ is the pooled replica-level standard deviation. With three treatment and three control replicas, residual *df* = 4 and *J* = 0.80.

Effects were oriented so that positive values consistently denoted greater restriction. Signs were reversed for conformational entropy, covariance trace, RMSF, and dihedral-transition count, for which decreases indicate restriction; the original sign was retained for effective stiffness, *S*^2^_min_, and persistence length. The aligned-metric count in Figure 2 is the number of positive oriented effects and was used descriptively rather than as a significance test. Additional details on metric orientation, the evidence-classification criteria, and the definition of contiguous residue-level candidate regions are provided in Supplementary Method S1.

**Figure 2.**
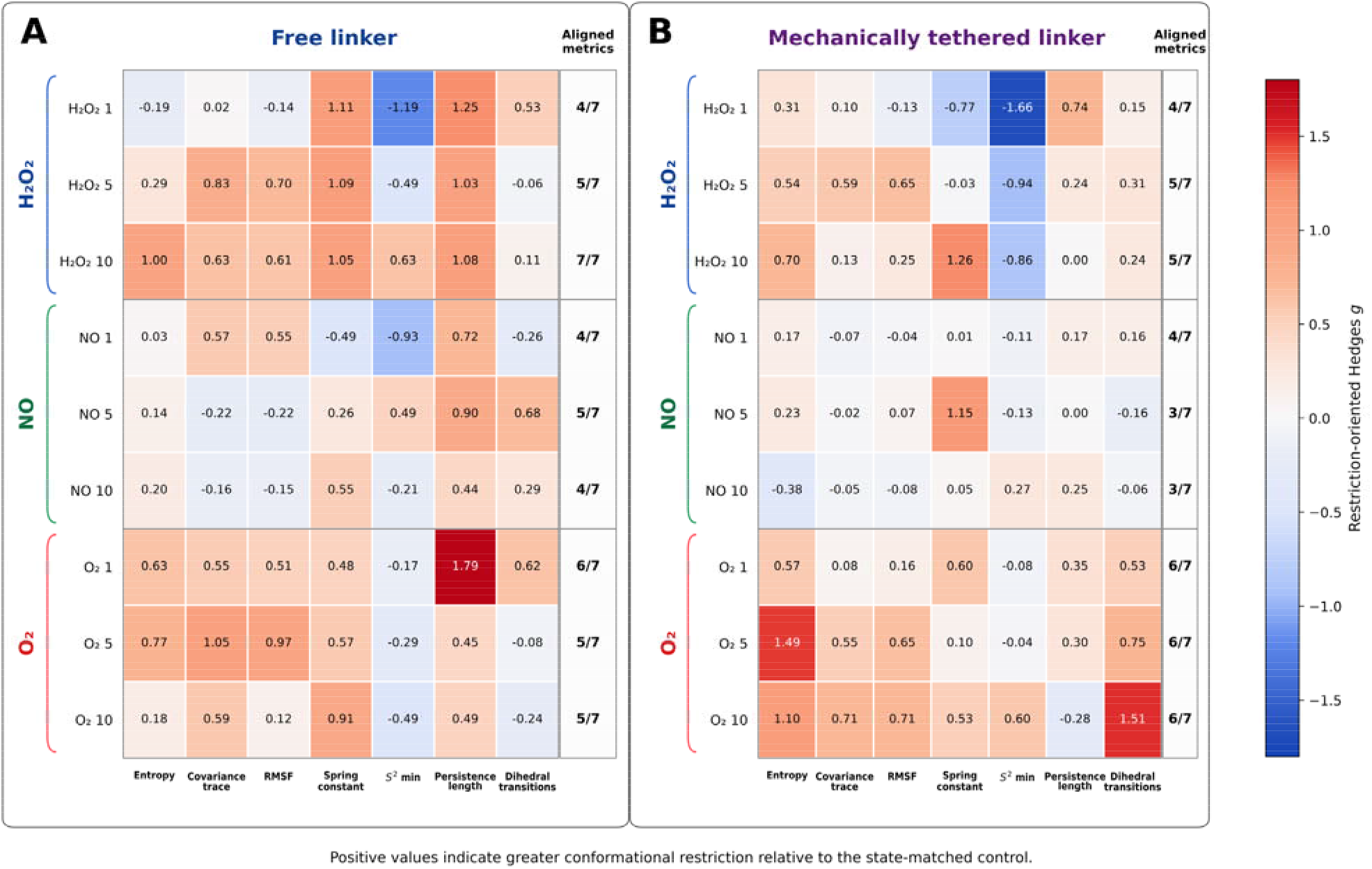
CAP-associated molecular effects are species-and loading-dependent in the free linker and remain detectable under terminal tethering. Restriction-oriented Hedges values are shown for treatment-versus-state-matched-control comparisons in FL (A) and TL (B). Positive values denote reduced entropy, covariance trace, RMSF, or dihedral exchange, or increased effective stiffness, backbone order, or persistence length. The aligned-metric column is a descriptive count of positive oriented effects. Panel A constitutes the permissive chemical screen; Panel B shows the same perturbations in the mechanically constrained follow-up model.

Modification by the mechanical boundary condition was quantified by the difference-in-differences interaction

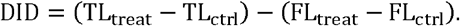

Positive oriented DID values indicate amplification of treatment-induced restriction by tethering; negative values indicate attenuation or reversal. The analysis included the six descriptors with commensurate FL and TL definitions: conformational entropy, covariance trace, RMSF, S_min_^2^, persistence length, and dihedral-transition count. The transformation of posterior DID draws to the restriction-oriented standardized scale is described in Supplementary Method S2. Complete descriptor-level Bayesian model diagnostics are provided in Source Data File.

Replica-level observations were modeled separately for each descriptor using a robust Bayesian factorial model with a Student-*t* likelihood (v = 4) and partial pooling across the nine species-by-loading interactions. Four chains were sampled, with 2600 retained posterior draws per chain after warm-up (approximately 10,400 draws per descriptor). Posterior raw interactions were transformed linearly to a restriction-oriented standardized scale using the four-cell residual variance (*df* = 8) and Hedges correction *J* = 0.9032. The posterior probability of amplification was

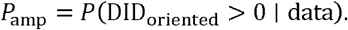

Directional evidence was classified as exploratory at P_amp_ ≥ 0.75, supported at P_amp_ ≥ 0.90, and strong at P_amp_ ≥ 0.95, provided that the direction remained stable in deletion analyses.

### 2.5. Projected conformational landscapes

Conformational landscapes were projected onto the joint radius-of-gyration–end-to-end-distance coordinate, (*R_g_*, r). A normalized 28 x 28 histogram was constructed for each replica using grid boundaries shared within each mechanical family. Replica distributions were averaged with equal weight and smoothed with a one-bin Gaussian kernel. Projected free energy was calculated as

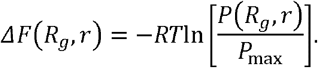

For visualization, adjacent 2 x 2 probability bins were aggregated to a 14 x14 grid and ΔF was recalculated; all quantitative statistics were derived from the original 28 x 28 grids. These surfaces are two-coordinate projected potentials of mean force, not complete molecular free-energy landscapes.

Compact, intermediate, and extended states were defined separately within FL and TL by applying *k*-means clustering (*k* = 3) to standardized (*R_g_*, *r*) values from the matched control trajectories only. Control-derived centroids were then fixed for treatment classification. State labels are therefore family-relative and do not imply identical absolute geometries in FL and TL.

State occupancies were calculated per replica. For switching analysis, coordinate trajectories were averaged in 100-ps blocks, assigned to the nearest control-defined centroid, and filtered with a three-block majority rule before transitions were counted and normalized per 10 ns. Treatment-control similarity was quantified by the probability-overlap coefficient,

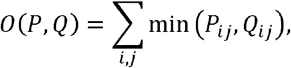

and by Jensen–Shannon divergence. Landscape breadth was summarized by the area within 2*RT* of the minimum and by the effective number of occupied bins. The replica-balanced bootstrap procedure, balanced deletion analyses, and early/late trajectory-window sensitivity tests used to evaluate the landscape contrasts are described in Supplementary Method S3.

### 2.6. Residue-level dynamics, hydrogen bonding, and interaction energetics

Residue-level restriction-oriented Hedges *g* profiles were calculated for Cα RMSF and backbone S_min_^2^. Interpretation focused on contiguous candidate regions rather than isolated residues. A region required at least three consecutive residues for which every residue satisfied | *g*| ≥ 0.80, posterior probability in the observed direction ≥ 0.80, and balanced replica-deletion stability ≥ 0.80. Regions were classified as exploratory, supported, or strong according to directional probability and deletion robustness.

Intramolecular linker hydrogen bonds were identified with the default GROMACS geometry: donor–acceptor distance ≤ 0.35 nm and H–D–A angular deviation ≤ 30, equivalent to a conventional D–H···A angle ≥ 150°. Counts were summarized per replica and expressed relative to the matched control.

Species–linker interaction energies were estimated with gmx_MMPBSA ^[46]^. Generalized Born calculations used the modified Onufriev–Bashford–Case model 2 (igb=5), 0.15 M ionic strength, internal and external dielectric constants of 1.0 and 78.5, respectively, and the default mbondi2 radii. The decomposition included van der Waals, electrostatic, polar-solvation, and nonpolar-solvation terms. Native outputs were multiplied by 1000 to obtain units of cal mol^-1^, and divided by the number of species molecules in each condition after replica averaging. Residue-level decompositions were compared with RMSF and S_min_^2^ profiles using rank-based alignment tests that preserved spatial autocorrelation; multiple comparisons were controlled within each energy–dynamics family. The resulting values were treated as comparative interaction-energy fingerprints; configurational entropy was not included.

A condition-level integrated restriction index was constructed from the six descriptors with comparable FL and TL definitions. For condition c and metric m, the raw treatment-control effect was restriction-oriented and standardized across the 18 single-species treatments,

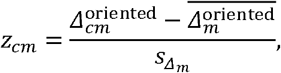

and the index was calculated as

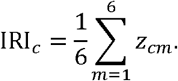

Spearman correlation assessed condition-level associations of the IRI with intramolecular H-bond change and net attractive MM/GBSA strength. Uncertainty was estimated by 10,000 nonparametric bootstrap resamples of the 18 treatment conditions. For energy–restriction analysis, a partial rank correlation was obtained by residualizing ranked energy and IRI values against log loading, species identity, and mechanical state. The circular-shift procedure used to preserve sequence autocorrelation and the prespecified energetic-colocalization criteria are described in Supplementary Method S4.

### 2.7. Robustness and sensitivity analyses

Key conclusions were evaluated by balanced leave-one-replica deletion and analysis-window sensitivity. Two-group landscape contrasts used the exact 729 ordered nonparametric bootstrap combinations; four-group boundary-condition contrasts used 200,000 bootstrap draws. Early-and late-window analyses tested the stability of occupancy, overlap, state-space breadth, modal coordinates, and switching. A result was described as replica robust only when its qualitative direction was retained under the relevant deletion and window analyses. Condition-specific landscape robustness results and trajectory-window sensitivity analyses are reported in Supplementary Table S4 and Supplementary Figure S2, respectively.

## 3. Results

### 3.1. CAP-associated molecular species remodel the unconstrained ABCB1 linker in species-and loading-dependent ways

We first asked whether the isolated ABCB1 linker was responsive to the selected CAP-associated molecular species. In the unconstrained FL model, H_2_O_2_, NO, and O_2_ produced distinct and nonmonotonic treatment–control phenotypes across the global descriptors (Figure 2A). All effects were evaluated relative to the FL control so that this first stage addressed chemical susceptibility within a single mechanical reference state.

H_2_O_2_ showed the clearest progressive FL response: four of seven restriction-oriented metrics were aligned at one molecule, five at five molecules, and all seven at ten molecules. NO produced a weaker whole-linker pattern, with small and nonmonotonic effects across loadings. O_2_ followed a different pattern: one molecule shifted six of seven FL metrics toward restriction, the response remained predominantly restrictive at five molecules, and became less coherent at ten. Thus, the screen established species-and loading-dependent chemical remodeling without a simple monotonic relation to molecular copy number.

The low-loading O_2_ condition provided the clearest ensemble-level example of this chemical susceptibility. FL + 1 O_2_ redistributed the projected ensemble toward a predominantly compact, low-exchange state and substantially reduced overlap with the FL control (Figure 3A,D). This strong response identified a chemically sensitive conformational route whose transferability could be tested by repeating the same perturbation under terminal tethering. Complete state-matched treatment effects, including control and treatment means, raw differences, restriction-oriented Hedges values, directional probabilities, and deletion-based robustness classifications, are provided in the Source Data File.

**Figure 3.**
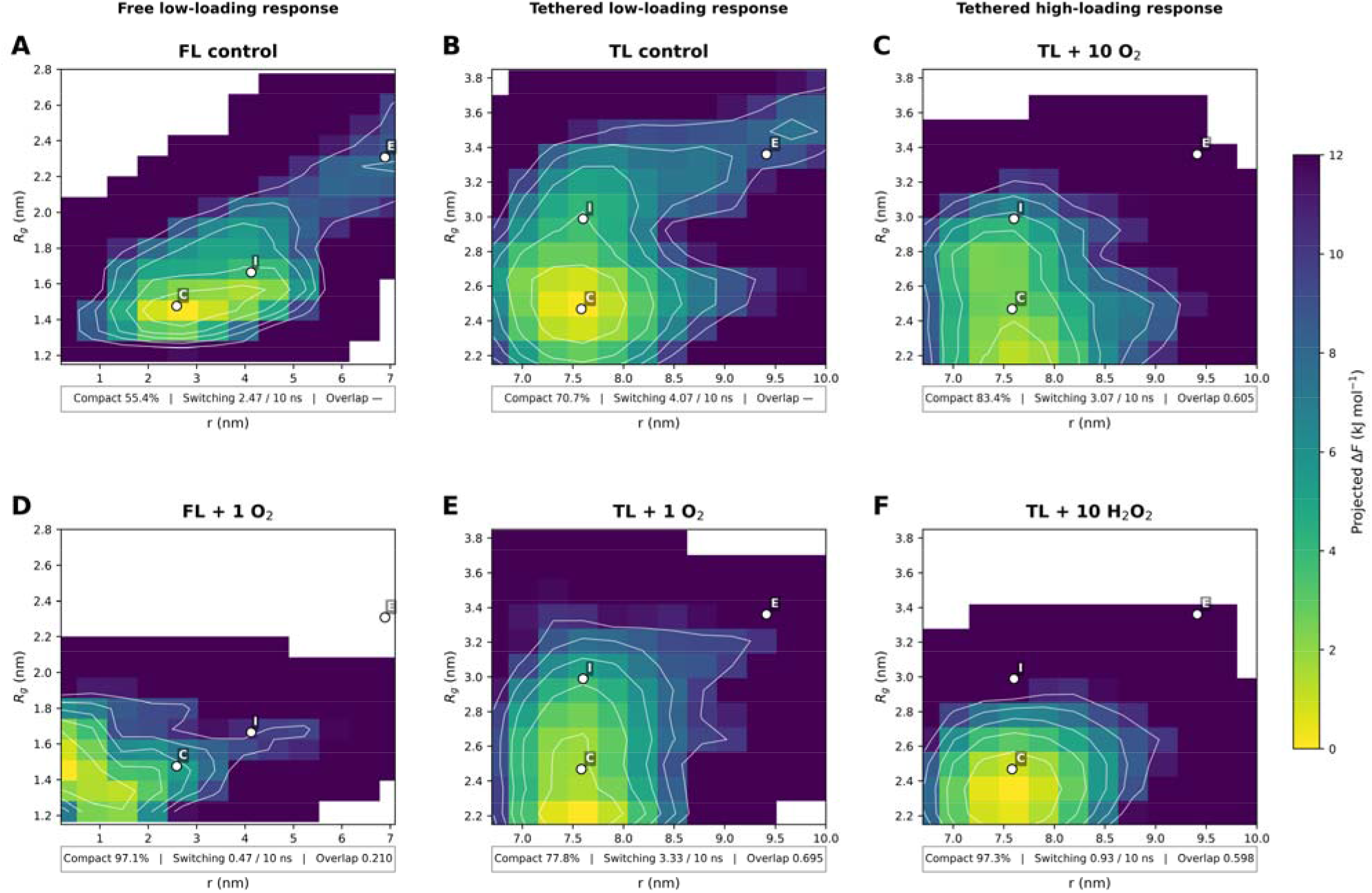
Mechanically constrained follow-up distinguishes retained from suppressed CAP-associated linker phenotypes. Replica-balanced projected free-energy landscapes are shown for FL control **(A)**, TL control **(B)**, TL + 10 OF **(C)**, FL + 1 OF **(D)**, TL + 1 OF **(E)**, and TL + 10 HF OF **(F)**. C, I, and E are compact, intermediate, and extended centroids defined from the matched control within each mechanical family. Summary strips report compact occupancy, switching per 10 ns, and overlap with the corresponding control.

### 3.2. Selected CAP-associated remodeling persists under structure-derived terminal tethering

We next asked whether chemical sensitivity persisted when terminal geometry was restricted to a structure-derived range. The unexposed FL and TL controls occupied different geometrical regimes, as expected from the terminal constraint, but TL was not uniformly more rigid: tethering maintained a larger terminal separation while preserving substantial internal fluctuations and conformational exchange. Consequently, every treatment was evaluated relative to its state-matched control, and FL and TL absolute state labels were not interpreted as directly equivalent geometries.

Against this TL reference, selected plasma-associated perturbations remained clearly detectable (Figure 2B). O_2_ effects were modest at one molecule but became broadly restrictive at ten, when six of seven metrics aligned and the strongest change involved reduced dihedral exchange (g = 1.51). H_2_O_2_ response remained more internally mixed, most notably because S_min_^2^ shifted toward lower local order even when global descriptors indicated restriction. Despite this mixed descriptor profile, high-loading H_2_O_2_ produced a coherent landscape response, with compact-state occupancy increasing from 70.7% to 97.3%, switching decreasing from 4.07 to 0.93 transitions per 10 ns, and the modal radius of gyration shifting from 2.555 to 2.342 nm (Figure 3F). High-loading O_2_ also retained a distinct phenotype: compact occupancy increased to 83.4%, the effective number of occupied bins decreased from 141.4 to 94.2, and overlap with the matched control decreased to 0.605 (Figure 3C). NO remained weaker at the whole-linker level, although localized dynamical responses were detected below. These results show that the structure-derived terminal constraint did not eliminate chemical responsiveness. The coexistence of persistent and suppressed phenotypes motivated a direct comparison of how tethering changed the expression of chemical responses.

### 3.3. Mechanical tethering filters rather than abolishes chemical responses

Direct comparison of the two reduced models separated responses that depended strongly on free terminal motion from those that remained accessible after tethering. Low-loading O_2_ provided the clearest contrast: the compact, low-exchange phenotype seen in FL was largely absent in the corresponding TL ensemble (Figure 3D,E). By contrast, distinct high-loading H_2_O_2_ and O_2_ phenotypes remained detectable in TL. The FL-only O_2_ response should therefore be interpreted as an intrinsic susceptibility of the isolated linker rather than as a direct prediction for intact ABCB1, whereas the mechanically persistent responses provide higher-priority candidates for full-length validation. Replica-level heterogeneity of the high-loading tethered O_2_ and H_2_O_2_ landscapes is shown in Supplementary Figure S1, and early/late trajectory-window analyses are shown in Supplementary Figure S2. Condition-level landscape statistics and replica-balanced robustness estimates are reported in Supplementary Tables S3 and S4, respectively.

Formal difference-in-differences analysis provided a complementary test of mechanical modification beyond the state-matched comparisons. H_2_O_2_ and NO showed no coherent species-wide amplification by tethering, whereas O_2_ developed a more positive directional pattern with loading. At ten O_2_ molecules, three of six commensurate descriptors crossed the directional-support threshold, with the strongest signal involving reduced dihedral exchange; however, none of the central 95% credible intervals excluded zero. The interaction analysis therefore supports selective, loading-dependent mechanical filtering rather than a uniform cross-boundary effect (Supplementary Figure S3 and Supplementary Table S2). Complete descriptor-level Bayesian diagnostics are provided in the Source Data File. Because whole-linker averages can conceal offsetting regional responses, we next mapped the remodeling along the sequence.

### 3.4 CAP-associated remodeling is spatially heterogeneous along the linker

Residue-level RMSF and backbone-order profiles showed that chemical remodeling was spatially heterogeneous rather than a uniform microscopic rigidification (Figure 4). In FL + 1 O_2_, RMSF restriction clustered around residues 648–652 and 654–659, whereas reduced local order appeared within part of the same central region and near residues 698–700. The compact, low-exchange ensemble therefore retained localized flexibility. The complete set of contiguous candidate regions underlying Figure 4 is provided in the Source Data File. Opposite-direction regions are included to avoid selective presentation, and the reported regions should not be interpreted as collections of independently significant residues.

**Figure 4.**
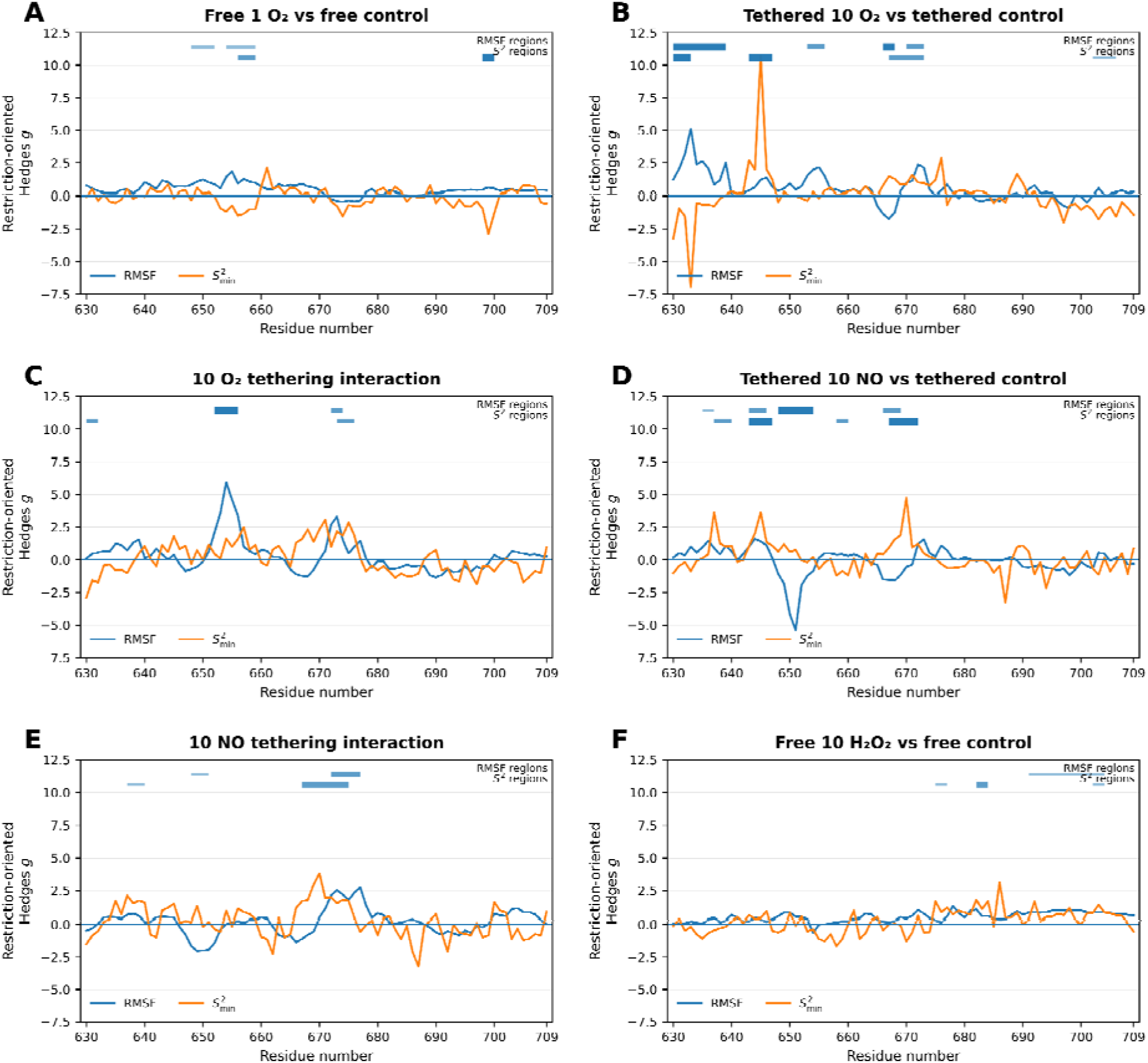
CAP-associated dynamical responses are spatially heterogeneous along the ABCB1 linker. Restriction-oriented residue-level Hedges g profiles are shown for RMSF and backbone s^2^. Positive values indicate local restriction in treatment-control panels and tethering-amplified restriction in DID panels. Horizontal tracks identify contiguous candidate regions satisfying the prespecified magnitude, directional-probability, and replica-deletion criteria. **(A)**, FL + 1 OF. **(B)**, TL + 10 OF. **(C)**, 10-OF tethering interaction. **(D)**, TL + 10 NO. (**E)**, 10-NO tethering interaction. **(F)**, FL + 10 HF OF.

High-loading TL O_2_ likewise produced discrete regional responses. The clearest RMSF restriction spanned residues 630–639, while increased backbone order occurred at 643–647 and additional central segments. The O_2_ interaction profile localized the strongest tethering-amplified RMSF restriction to residues 652–656, with a second overlapping RMSF–order response near 672–676. Opposite-direction shifts elsewhere show that a retained global phenotype can coexist with regional flexibility and that mechanical embedding redistributes, rather than simply clamps, local dynamics.

NO further illustrates why whole-linker averages can obscure chemical susceptibility. Although its global cross-boundary effects were near zero, TL + 10 NO produced localized ordering alongside increased RMSF in neighboring segments, while FL + 10 H_2_O_2_ showed a distinct C-terminal signature. Across species, contiguous replica-robust regions—not isolated residue peaks—defined the local response. This spatial heterogeneity prompted us to ask whether remodeling followed direct species–linker energetic attraction or instead tracked reorganization within the linker itself.

### 3.5. Internal linker-network reorganization better tracks remodeling than total species–linker attraction

MM/GBSA decomposition confirmed that the three molecular species retained distinct average energetic fingerprints (Figure 5A). H_2_O_2_ was dominated by electrostatic attraction that was substantially offset by polar-solvation penalty: in FL, mean per-molecule values were approximately 114.5 and 106.1 cal molO¹, yielding a net attractive strength of 28.9 cal molO¹; corresponding TL values were 85.8, 86.8, and 18.7 cal molO¹. NO and O_2_ were predominantly van der Waals driven, with net attractions of 11.8 and 4.7 cal molO¹ in FL and 15.0 and 5.1 cal molO¹ in TL, respectively.

**Figure 5.**
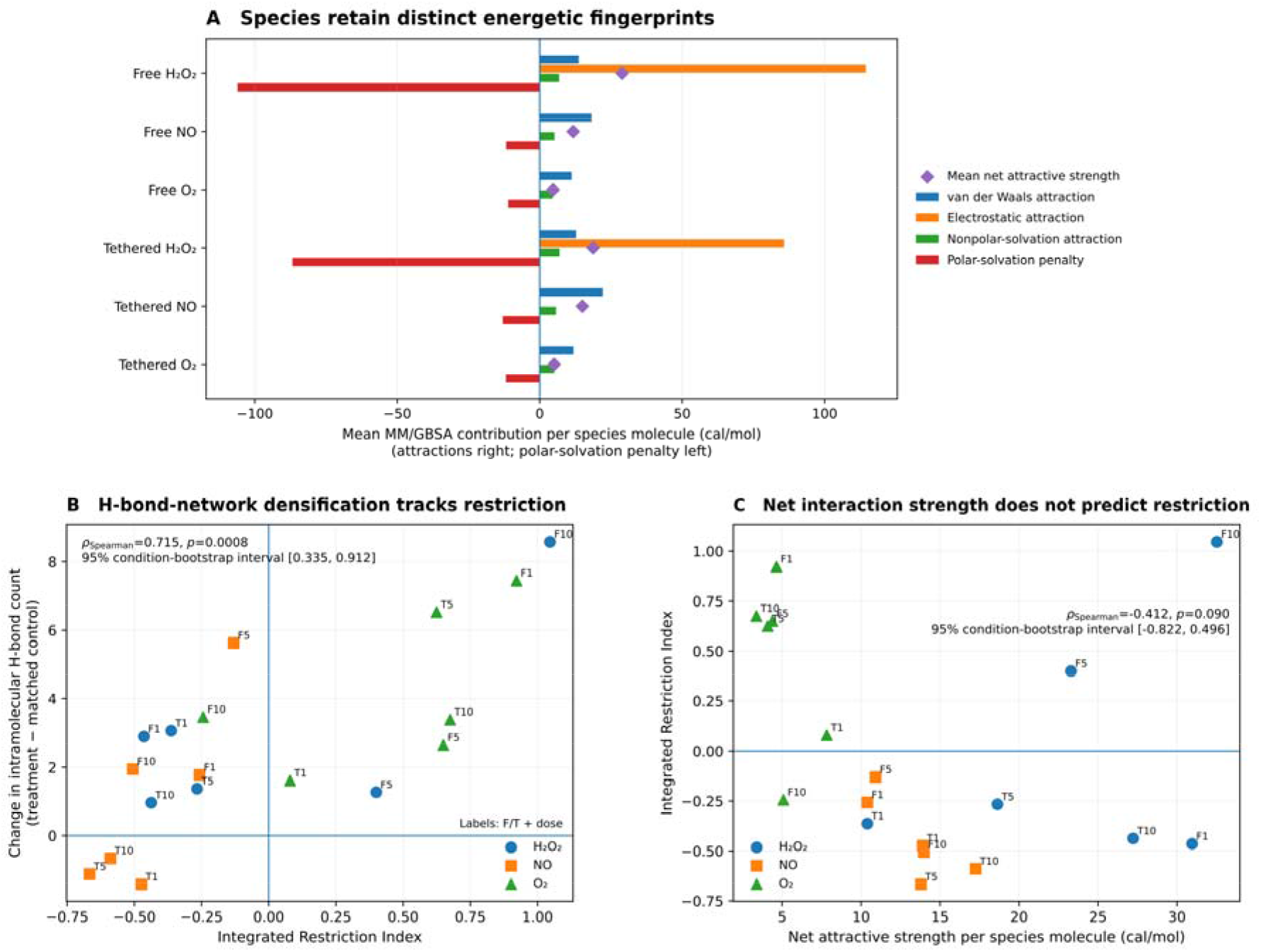
Interaction-energy fingerprints and intramolecular H-bond-network changes show different relationships with the integrated restriction index. (A), Mean per-molecule MM/GBSA energetic fingerprints for HFOF, NO, and OF in FL and TL, averaged across loadings and replicas. Values are reported in cal mol^-1^ per species molecule. (B), Association between the integrated restriction index and treatment-induced change in intramolecular H-bond count across 18 treatment conditions. (C), Net attractive strength versus the integrated restriction index. F and T denote free and tethered models, respectively; the number denotes molecular loading.

Total energetic attraction did not reproduce the ordering of conformational responses. O_2_ had the weakest net attraction yet generated the most pronounced low-loading FL redistribution, whereas the strongly polar H_2_O_2_ interaction did not consistently yield the largest conformational response. Net attraction was not reliably associated with the integrated restriction index (Figure 5C), either before or after adjustment for species, loading, and mechanical state. Average comparative MM/GBSA attraction therefore did not provide a sufficient predictor of ensemble remodeling.

Intramolecular H-bond reorganization tracked restriction more closely (Figure 5B). Treatment-induced H-bond-count change correlated positively with the integrated restriction index, with a 95% bootstrap interval of 0.335–0.912. This association does not establish causality, but it identifies internal network reorganization as a closer structural correlate of restriction than total species–linker attraction. Residue-level energetic attraction likewise failed to provide a universal predictor of local rigidification. Prespecified regional energetic alignments meeting the colocalization criteria are listed in Supplementary Table S6, and complete condition-level inputs are provided in the Source Data File.

## 4. Discussion

### 4.1. Chemical remodeling reflects ensemble redistribution rather than simple dose or interaction strength

The central mechanistic implication is that the ABCB1 linker behaves as an ensemble-responsive intrinsically disordered segment rather than as a site that must undergo stable ligand binding to be perturbed. The nonmonotonic species-and loading-dependent patterns argue against treating finite molecular copy number as a conventional linear dose axis. A more consistent interpretation is that transient contacts bias pre-existing conformational basins, with the observed response depending jointly on chemical identity and the states that are accessible when a contact occurs.

The contrasting H_2_O_2_ and O_2_ behaviors illustrate two possible routes to such redistribution. A progressively coherent H_2_O_2_ phenotype is compatible with distributed weak perturbations becoming collectively expressed across the ensemble, whereas the disproportionate low-loading O_2_ response suggests that a single molecule can have a large effect when it stabilizes a mechanically accessible state. These interpretations do not establish a unique microscopic pathway, but they explain why increasing copy number did not simply scale the same conformational phenotype.

The energetic analysis reinforces this ensemble-based view. Comparative MM/GBSA attraction did not rank conditions by their conformational response, indicating that cumulative attraction alone is insufficient to predict remodeling. For an intrinsically disordered region, where contacts are transient and multiple conformations can be close in free energy, the location and timing of interactions may matter more than their time-averaged total strength. This distinction is important for CAP-associated chemistry because it separates chemical contact strength from mechanical consequence.

### 4.2. Mechanical tethering functions as a transferability filter

The value of the tethered model is therefore not that it creates a generically more rigid linker, but that it changes the question from whether the isolated chain can respond to whether a chemical phenotype survives restricted terminal geometry. The TL control remained dynamically active, showing that loss of a phenotype under tethering cannot be attributed simply to immobilization. Instead, the restraint changes the subset of conformational routes available to the perturbation.

This distinction separates two classes of response. Phenotypes seen only in FL reveal intrinsic chemical susceptibility that depends strongly on unconstrained terminal motion, whereas responses retained in TL remain accessible within a structure-derived mechanical envelope. The suppressed low-loading O_2_ transition and the persistence of selected high-loading H_2_O_2_ and O_2_ phenotypes therefore serve different interpretive roles: the former defines a free-motion-dependent route, while the latter identify mechanically persistent candidates for subsequent full-length testing.

The formal boundary-condition analysis supports this selective interpretation rather than a global gating model. Broad credible intervals and mixed descriptor-level interactions do not justify claiming that tethering uniformly amplifies or attenuates chemical effects. The principal value of the FL–TL comparison is instead discriminatory: together, the two reduced models identify which responses are especially sensitive to mechanical embedding and which remain plausible after terminal freedom is restricted.

### 4.3. Regional and intramolecular network reorganization underlie the global phenotypes

Global labels such as compaction or restriction should not be interpreted as chain-wide rigidification. The residue-level profiles show that locally restricted and locally flexible segments can coexist within the same ensemble. Such opposition is expected for an intrinsically disordered sequence and provides a natural explanation for conditions in which whole-linker averages are modest despite reproducible local responses.

For this reason, contiguous response regions are more informative than isolated residue peaks. Neighboring fluctuations are correlated, and local changes may redistribute transient contacts or backbone preferences without creating a single rigid state. The candidate regions identified here should therefore be viewed as hypotheses for where linker remodeling might couple to the surrounding transporter, not as collections of independently significant residues.

Intramolecular hydrogen-bond reorganization provides a complementary structural correlate of this regional remodeling. Its closer association with the integrated restriction index, together with the poor predictive value of total species–linker attraction, is consistent with chemical perturbations being transduced through reorganization of the linker’s internal contact network. The direction of causality remains unresolved: additional hydrogen bonds may help stabilize restricted ensembles, or they may arise because compaction increases opportunities for internal contact.

### 4.4. Potential consequences for ABCB1 interdomain coordination and drug efflux

Taken together, the simulations support a constrained mechanistic hypothesis linking CAP-associated chemistry to P-gp regulation. CAP or plasma-activated media can enhance chemotherapy responses in resistant cancer models [30–34], and the ABCB1 linker requires structural flexibility for efficient ATP hydrolysis and drug transport [17,18]. The present work connects these observations at one intermediate step by showing that selected plasma-associated molecules can remodel the linker ensemble and that some responses remain detectable after terminal geometry is mechanically restricted.

The downstream consequences remain untested. The reduced models do not establish altered NBD–TMD communication, ATPase activity, substrate transport, or multidrug-resistance reversal. A useful next step is therefore to test propagation rather than simply reproduce the reduced-linker metrics: full-length, membrane-embedded ABCB1 simulations should ask whether mechanically persistent linker perturbations alter NBD separation, coupling-helix or intracellular-loop interactions, nucleotide-dependent conformational transitions, or substrate-access pathways.

Functional experiments can then determine whether those structural perturbations have transport consequences. ATPase activity, fluorescent-substrate efflux, and intracellular chemotherapeutic accumulation could be measured after controlled exposure to plasma-activated media or defined molecular-species conditions. Linker mutants or constructs with altered flexibility would provide a direct test of linker dependence, while the contiguous candidate regions identified here offer initial targets for perturbation. Such validation is required before linker remodeling can be connected causally to drug-efflux modulation in cells.

### 4.5. Implications and limitations

Several limitations bound the interpretation. The simulated construct is an isolated, solvated peptide corresponding to the P-gp linker sequence rather than a reconstruction of the linker within full-length ABCB1. TL is likewise a mechanically constrained reduced representation, not a physiological model: it restores a structure-derived terminal-separation range but does not reproduce terminal orientation, direct contacts with the surrounding domains and intracellular loops, or the membrane environment. The isolated constructs also contain ionized N-and C-terminal groups at residues 628 and 711, whereas these positions are peptide-bonded in full-length ABCB1; terminal electrostatics may therefore influence the accessible ensemble, particularly in FL.

Sampling and model chemistry impose additional limits. The simulations began from one AlphaFold2 conformation, and three 100-ns replicas cannot exhaust the ensemble of an intrinsically disordered chain. The projected landscapes are reduced-coordinate descriptions rather than validated Markov-state models, and the broad difference-in-differences credible intervals reflect the small number of independent replicas. MM/GBSA omits configurational entropy and was used only as a comparative interaction-energy fingerprint. Classical MD captures the modeled noncovalent interactions but not oxidation, radical chemistry, or other covalent modifications that may accompany plasma exposure. Finally, the molecule counts are operational simulation loadings rather than direct proxies for biological concentrations, and mixed-species effects are outside the scope of the present study.

### 4.6. Conclusion

Selected CAP-associated molecular species altered the conformational ensemble of the ABCB1 NBD1–TMD2 linker through species-and loading-dependent noncovalent effects. The two-stage design separated chemical susceptibility from mechanical transferability: some responses depended strongly on free terminal motion, whereas distinct H_2_O_2_-and O_2_-associated phenotypes remained detectable under structure-derived tethering. These mechanically persistent responses do not demonstrate altered P-gp transport, but they provide concrete candidates for full-length simulations and functional tests of whether linker remodeling can propagate to NBD–TMD coordination, ATPase–transport coupling, and drug efflux.

## Declarations

### CRediT

Mansoureh Hoseini: Conceptualization, Methodology, Software, Formal analysis, Investigation, Data curation, Visualization, Writing – original draft. Hossein Abbasi: Supervision, Conceptualization, Methodology, Writing – review & editing.

### Competing interests

The authors declare no competing financial interests or personal relationships that could influence this work.

### Funding

This research received no specific grant from funding agencies in the public, commercial, or not-for-profit sectors.

### Data availability

All replica-level results are in Paper1_Source_Data.csv (supplementary). Input topologies/analysis scripts available from the corresponding author on reasonable request.

## Supporting information

Paper1_Source_Data

## Supplementary Information

### Supplementary Methods

#### Supplementary Method S1. Restriction orientation and evidence categories

Treatment-control effects were calculated from replica-level means. Metric signs were oriented so that positive values represented greater restriction: entropy, covariance trace, RMSF, and dihedral-transition count were multiplied by-1, whereas effective stiffness, S2min, and persistence length retained their original sign. The aligned-metric count in Figure 2 is descriptive and was not used as an inferential test.

For residue-level profiles, candidate regions required at least three consecutive residues with |g| >= 0.80, directional probability >= 0.80, and balanced deletion stability >= 0.80. Regions were classified as exploratory, supported, or strong according to the prespecified probability and robustness rules used in the main analysis.

#### Supplementary Method S2. Posterior DID standardization

The raw interaction was defined as (TL treatment - TL control) - (FL treatment - FL control). Posterior draws were transformed linearly to the restriction-oriented standardized scale using the corresponding pooled-variance factor, the metric-specific orientation, residual df = 8, and Hedges correction J = 0.9032. P_amp was the fraction of oriented posterior draws above zero. The Student-t likelihood parameter nu = 4 is distinct from the residual degrees of freedom used for standardization.

#### Supplementary Method S3. Landscape robustness

The principal landscape contrasts were evaluated using replica-balanced probabilities. Two-group contrasts used the exact set of 729 ordered bootstrap combinations, and four-group boundary-condition contrasts used 200,000 bootstrap draws. Balanced leave-one-replica analyses and early/late trajectory windows were used to evaluate directional stability. The reduced-coordinate states are control-defined family-relative states and do not constitute a Markov-state model.

#### Supplementary Method S4. Energy-dynamics alignment

Residue-level energy-dynamics associations were assessed using Spearman rank correlation and circular shifts along the sequence to preserve spatial autocorrelation. Multiplicity was controlled within each comparison family. Energetic colocalization was reported only when a candidate dynamical region overlapped a strong attractive-energy window under the prespecified alignment rules. MM/GBSA values are comparative interaction fingerprints, not absolute binding free energies.

The complete simulation protocol and definitions of the primary conformational descriptors are provided in the main Materials and Methods. The Supplementary Methods therefore focus only on analysis details needed to interpret the supporting tables, robustness checks, and boundary-condition comparisons.

## Supplementary Tables

**Table S1.** Replica-level baseline descriptors for the FL and TL controls.

| Descriptor | FL R1 | FL R2 | FL R3 | FL mean | FL SD | TL R1 | TL R2 | TL R3 | TL mean | TL SD | Unit |
| --- | --- | --- | --- | --- | --- | --- | --- | --- | --- | --- | --- |
| Conformational entropy | 3481.3 | 3327.3 | 3086.3 | 3298.3 | 199.10 | 3906.6 | 4058.1 | 3797.7 | 3920.8 | 130.75 | J mol<sup>-1</sup> K<sup>-1</sup> |
| Covariance trace | 224.60 | 176.71 | 36.616 | 145.97 | 97.687 | 163.38 | 251.14 | 155.75 | 190.09 | 53.007 | nm<sup>2</sup> |
| Mean C-alpha RMSF | 0.8620 | 0.7699 | 0.3533 | 0.6618 | 0.2710 | 0.7571 | 0.9722 | 0.7553 | 0.8282 | 0.1247 | nm |
| Effective stiffness | 5.1610 | 11.582 | 32.595 | 16.446 | 14.349 | 11.7955 | 6.6627 | 20.0325 | 12.8302 | 6.7447 | kJ mol<sup>-1</sup> nm<sup>-2</sup> |
| Minimum backbone order parameter | 0.7590 | 0.7314 | 0.7770 | 0.7558 | 0.0230 | 0.6611 | 0.7539 | 0.6862 | 0.7004 | 0.0480 | dimensionless |
| Persistence length | 0.9600 | 1.0200 | 0.9400 | 0.9733 | 0.0416 | 0.9900 | 0.9200 | 1.0400 | 0.9833 | 0.0603 | backbone-bond units |
| Dihedral-transition count | 3818.0 | 5279.0 | 5874.0 | 4990.3 | 1058.0 | 6399.0 | 6516.0 | 5010.0 | 5975.0 | 837.76 | count / 50 ns |
| Radius of gyration | 1.8223 | 1.5639 | 1.4446 | 1.6103 | 0.1931 | 2.3950 | 2.8580 | 2.4720 | 2.5750 | 0.2481 | nm |
| End-to-end distance | 3.8647 | 3.9149 | 2.6778 | 3.4858 | 0.7002 | 7.7900 | 7.7420 | 7.5880 | 7.7067 | 0.1055 | nm |
| Intramolecular H-bond count | 39.010 | 46.400 | 40.820 | 42.077 | 3.8519 | 43.068 | 34.302 | 43.221 | 40.197 | 5.1058 | count |

Values are individual replica summaries followed by the mean and between-replica SD. FL and TL effective stiffness estimates were obtained using different mechanical definitions and were not compared directly.

Complete state-matched treatment effects underlying ***Figure 2*** are provided in Source_Data_File_1_State_Matched_Effects data set in Source Data File. The file contains all 126 treatment-by-descriptor rows, including control and treatment means, raw differences, restriction-oriented Hedges g values, directional probabilities, leave-one-out sign stability, robustness classifications, and units.

**Table S2.** Complete posterior boundary-condition interactions underlying Supplementary Figure S3.

| Species | Dose | Short_Name | DID | Standardized_DID_g | Posterior standardized mean | 95% CrI low | 95% CrI high | P_Amplification | Single_Delete_Sign_Stable | Balanced_Delete_Sign_Fraction | Robustness | Amplification_Support |
| --- | --- | --- | --- | --- | --- | --- | --- | --- | --- | --- | --- | --- |
| H2O2 | 1 | CovTrace | -7.1894 | -0.0859 | 0.0440 | -0.7310 | 0.8746 | 0.5374 | No | 0.531 | Uncertain | No directional support |
| H2O2 | 1 | DihedralCount | 446.00 | 0.4900 | -0.1446 | -1.1065 | 0.7779 | 0.3760 | No | 0.716 | Uncertain | No directional support |
| H2O2 | 1 | Entropy | -221.39 | -0.5718 | 0.0861 | -0.8921 | 1.0790 | 0.5645 | No | 0.790 | Uncertain | No directional support |
| H2O2 | 1 | Persistence | -0.0033 | -0.0624 | 0.2780 | -0.5322 | 1.1453 | 0.7438 | No | 0.519 | Uncertain | No directional support |
| H2O2 | 1 | RMSF | -0.0181 | -0.0915 | 0.0795 | -0.8400 | 0.9764 | 0.5677 | No | 0.617 | Uncertain | No directional support |
| H2O2 | 1 | S2Min | -0.0075 | -0.1652 | -0.4004 | -1.7724 | 0.8940 | 0.2715 | No | 0.568 | Uncertain | No directional support |
| H2O2 | 5 | CovTrace | 41.854 | 0.6073 | 0.0312 | -0.8895 | 0.9479 | 0.5267 | Yes | 0.765 | Uncertain | No directional support |
| H2O2 | 5 | DihedralCount | -395.00 | -0.4015 | 0.1465 | -0.6935 | 1.0446 | 0.6315 | No | 0.704 | Uncertain | No directional support |
| H2O2 | 5 | Entropy | -36.980 | -0.1456 | 0.0871 | -1.3183 | 1.4940 | 0.5511 | No | 0.580 | Uncertain | No directional support |
| H2O2 | 5 | Persistence | -0.0533 | -1.0045 | -0.0526 | -0.8531 | 0.7374 | 0.4513 | Yes | 0.914 | Uncertain | No directional support |
| H2O2 | 5 | RMSF | 0.0897 | 0.4637 | 0.0437 | -0.8813 | 0.9860 | 0.5321 | No | 0.691 | Uncertain | No directional support |
| H2O2 | 5 | S2Min | -0.0380 | -0.5266 | -0.4005 | -1.3862 | 0.5298 | 0.2008 | Yes | 0.728 | Exploratory | Weak attenuation/reversal support |
| H2O2 | 10 | CovTrace | 55.368 | 0.7237 | -0.0736 | -0.9209 | 0.7529 | 0.4327 | Yes | 0.815 | Uncertain | No directional support |
| H2O2 | 10 | DihedralCount | -90.000 | -0.1002 | 0.0100 | -0.8856 | 0.9392 | 0.5095 | No | 0.593 | Uncertain | No directional support |
| H2O2 | 10 | Entropy | 212.84 | 0.7788 | -0.1224 | -1.4427 | 1.2011 | 0.4251 | Yes | 0.852 | Uncertain | No directional support |
| H2O2 | 10 | Persistence | -0.0700 | -1.2752 | -0.1588 | -0.9597 | 0.6001 | 0.3434 | Yes | 0.975 | Uncertain | No directional support |
| H2O2 | 10 | RMSF | 0.1524 | 0.7029 | -0.0750 | -0.9251 | 0.7600 | 0.4259 | Yes | 0.827 | Uncertain | No directional support |
| H2O2 | 10 | S2Min | -0.0680 | -1.6996 | -0.6950 | -2.2841 | 0.8055 | 0.1853 | Yes | 0.988 | Exploratory | Weak attenuation/reversal support |
| NO | 1 | CovTrace | 69.015 | 0.8513 | -0.1552 | -0.9638 | 0.6503 | 0.3497 | Yes | 0.901 | Uncertain | No directional support |
| NO | 1 | DihedralCount | -475.33 | -0.4855 | 0.1163 | -0.7578 | 1.0196 | 0.5964 | No | 0.753 | Uncertain | No directional support |
| NO | 1 | Entropy | -39.553 | -0.0967 | -0.0501 | -0.9726 | 0.8591 | 0.4574 | No | 0.593 | Uncertain | No directional support |
| NO | 1 | Persistence | -0.0200 | -0.3471 | 0.1362 | -0.6590 | 0.9275 | 0.6419 | No | 0.642 | Uncertain | No directional support |
| NO | 1 | RMSF | 0.1895 | 0.8241 | -0.1827 | -1.0207 | 0.6019 | 0.3287 | Yes | 0.877 | Uncertain | No directional support |
| NO | 1 | S2Min | 0.0740 | 0.9196 | 0.0561 | -0.7203 | 0.8918 | 0.5479 | Yes | 0.889 | Uncertain | No directional support |
| NO | 5 | CovTrace | -26.425 | -0.2930 | 0.1294 | -0.6137 | 0.9168 | 0.6324 | No | 0.654 | Uncertain | No directional support |
| NO | 5 | DihedralCount | 1082.3 | 0.9675 | -0.3467 | -1.2126 | 0.4872 | 0.2084 | Yes | 0.914 | Exploratory | Weak attenuation/reversal support |
| NO | 5 | Entropy | 15.750 | 0.0535 | 0.0225 | -1.1881 | 1.2306 | 0.5117 | No | 0.568 | Uncertain | No directional support |
| NO | 5 | Persistence | -0.0367 | -0.6906 | -0.1145 | -0.9636 | 0.7158 | 0.3895 | Yes | 0.815 | Uncertain | No directional support |
| NO | 5 | RMSF | -0.0855 | -0.3617 | 0.1603 | -0.6036 | 0.9569 | 0.6611 | No | 0.704 | Uncertain | No directional support |
| NO | 5 | S2Min | -0.0310 | -0.6030 | -0.1061 | -1.3451 | 1.0806 | 0.4382 | No | 0.778 | Uncertain | No directional support |
| NO | 10 | CovTrace | -23.431 | -0.2283 | 0.0110 | -0.5948 | 0.6438 | 0.5101 | No | 0.593 | Uncertain | No directional support |
| NO | 10 | DihedralCount | 495.00 | 0.4284 | -0.1470 | -0.9128 | 0.6075 | 0.3484 | No | 0.704 | Uncertain | No directional support |
| NO | 10 | Entropy | 204.91 | 0.5961 | -0.1522 | -1.2457 | 0.9100 | 0.3897 | No | 0.790 | Uncertain | No directional support |
| NO | 10 | Persistence | -0.0033 | -0.0657 | 0.1710 | -0.6867 | 1.0358 | 0.6594 | No | 0.519 | Uncertain | No directional support |
| NO | 10 | RMSF | -0.0471 | -0.1847 | 0.0276 | -0.6880 | 0.7434 | 0.5332 | No | 0.556 | Uncertain | No directional support |
| NO | 10 | S2Min | 0.0405 | 0.5405 | 0.2172 | -0.6537 | 1.1266 | 0.6923 | No | 0.790 | Uncertain | No directional support |
| O2 | 1 | CovTrace | 54.041 | 0.5640 | -0.0722 | -0.7991 | 0.6435 | 0.4168 | Yes | 0.741 | Uncertain | No directional support |
| O2 | 1 | DihedralCount | 244.33 | 0.2943 | -0.0328 | -0.9931 | 0.9348 | 0.4720 | No | 0.642 | Uncertain | No directional support |
| O2 | 1 | Entropy | 36.567 | 0.1074 | 0.0154 | -1.0865 | 1.1443 | 0.5076 | No | 0.543 | Uncertain | No directional support |
| O2 | 1 | Persistence | -0.0400 | -0.6699 | -0.0632 | -0.8154 | 0.7121 | 0.4299 | Yes | 0.790 | Uncertain | No directional support |
| O2 | 1 | RMSF | 0.1286 | 0.4832 | -0.0723 | -0.8252 | 0.6612 | 0.4234 | Yes | 0.741 | Uncertain | No directional support |
| O2 | 1 | S2Min | 0.0043 | 0.1025 | 0.1236 | -1.2890 | 1.5047 | 0.5738 | No | 0.556 | Uncertain | No directional support |
| O2 | 5 | CovTrace | 53.560 | 0.7274 | 0.0320 | -0.8384 | 0.9196 | 0.5293 | Yes | 0.827 | Uncertain | No directional support |
| O2 | 5 | DihedralCount | -866.67 | -0.9092 | 0.2975 | -0.5556 | 1.2506 | 0.7462 | Yes | 0.840 | Uncertain | No directional support |
| O2 | 5 | Entropy | -47.503 | -0.1608 | 0.2855 | -0.9399 | 1.5272 | 0.6808 | No | 0.593 | Uncertain | No directional support |
| O2 | 5 | Persistence | -0.0000 | -0.0000 | 0.0570 | -0.6886 | 0.8207 | 0.5573 | No | 0.469 | Uncertain | No directional support |
| O2 | 5 | RMSF | 0.1420 | 0.7041 | -0.0076 | -0.9337 | 0.9122 | 0.4951 | Yes | 0.815 | Uncertain | No directional support |
| O2 | 5 | S2Min | 0.0178 | 0.2894 | 0.1028 | -0.9132 | 1.1237 | 0.5811 | No | 0.667 | Uncertain | No directional support |
| O2 | 10 | CovTrace | 16.956 | 0.2571 | 0.1608 | -0.7551 | 1.1229 | 0.6265 | No | 0.568 | Uncertain | No directional support |
| O2 | 10 | DihedralCount | -1621.0 | -1.6735 | 0.6597 | -0.2128 | 1.6541 | 0.9277 | Yes | 0.975 | Supported | Moderate amplification support |
| O2 | 10 | Entropy | -265.25 | -0.8792 | 0.4379 | -0.7503 | 1.6776 | 0.7651 | Yes | 0.877 | Exploratory | Weak amplification support |
| O2 | 10 | Persistence | -0.0467 | -0.8227 | -0.1695 | -0.9712 | 0.6085 | 0.3297 | Yes | 0.852 | Uncertain | No directional support |
| O2 | 10 | RMSF | -0.0553 | -0.2485 | 0.1538 | -0.6416 | 0.9677 | 0.6471 | No | 0.642 | Uncertain | No directional support |
| O2 | 10 | S2Min | 0.0651 | 1.2058 | 0.4996 | -0.6285 | 1.7221 | 0.8037 | Yes | 0.926 | Exploratory | Weak amplification support |

Positive posterior standardized values indicate amplification of restriction by tethering. Effective stiffness was excluded because the FL and TL estimators were not commensurate. CrI, central posterior credible interval.

*Source Data File. Complete descriptor-level Bayesian model diagnostics are provided in Source_Data_File_2_Model_Diagnostics. Data set in Source Data File*.

All 32 reported models passed the prespecified convergence criteria. The largest reported R-hat was 1.0007 for the FL treatment models, 1.0011 for the TL treatment models, and 1.0027 for the DID interaction models. Treatment models used four chains with 2,400 retained draws per chain; DID models used four chains with 2,600 retained draws per chain. The Student-t likelihood used nu = 4 throughout.

**Table S3.**
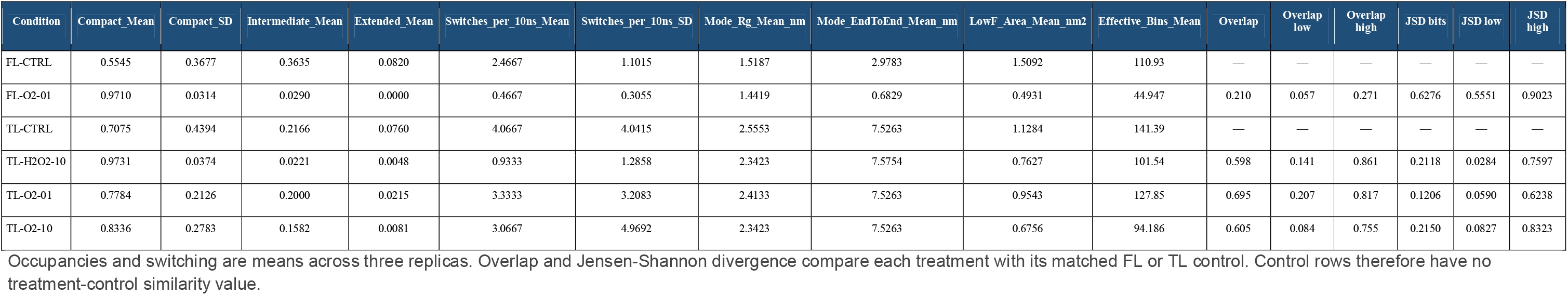
Condition-level landscape statistics for the six ensembles displayed in main Figure 3.

**Table S4.** Robustness of the landscape contrasts used in the main Results.

| Condition | Metric | Control_Mean | Treatment_Mean | Observed_Contrast | Bootstrap_2.5% | Bootstrap_97.5% | P_Observed_Direction | LOO_Sign_Stability | Window_Sign_Stable | Classification | Unit |
| --- | --- | --- | --- | --- | --- | --- | --- | --- | --- | --- | --- |
| FL-O2-01 | compact_occupancy | 0.5545 | 0.9710 | 0.4165 | 0.0075 | 0.7027 | 0.9904 | 1.000 | 1.0000 | Strong | fraction |
| FL-O2-01 | switches_per_10ns | 2.4667 | 0.4667 | -2.0000 | -2.8667 | -0.8000 | 1.0000 | 1.000 | — | Strong | transitions per 10 ns |
| FL-O2-01 | lowF_area_nm2 | 1.5092 | 0.4931 | -1.0162 | -2.7659 | 0.0180 | 0.9636 | 1.000 | — | Strong | nm^2 |
| FL-O2-01 | effective_bins | 110.93 | 44.947 | -65.987 | -182.12 | 8.7242 | 0.9588 | 1.000 | — | Strong | effective bins |
| FL-O2-01 | mode_rg_nm | 1.5187 | 1.4419 | -0.0768 | -0.2305 | 0.0768 | 0.8230 | 0.778 | — | Exploratory | nm |
| TL-H2O2-10 | compact_occupancy | 0.7075 | 0.9731 | 0.2657 | -0.0178 | 0.7718 | 0.9150 | 1.000 | 1.0000 | Supported | fraction |
| TL-H2O2-10 | switches_per_10ns | 4.0667 | 0.9333 | -3.1333 | -7.3333 | 0.5067 | 0.9424 | 1.000 | — | Supported | transitions per 10 ns |
| TL-H2O2-10 | lowF_area_nm2 | 1.1284 | 0.7627 | -0.3657 | -1.1493 | 0.1358 | 0.8107 | 0.667 | — | Exploratory | nm^2 |
| TL-H2O2-10 | effective_bins | 141.39 | 101.54 | -39.858 | -111.42 | 3.2700 | 0.9232 | 0.889 | — | Supported | effective bins |
| TL-H2O2-10 | mode_rg_nm | 2.5553 | 2.3423 | -0.2130 | -0.4022 | -0.0237 | 0.9863 | 1.000 | — | Strong | nm |
| TL-O2-01 | compact_occupancy | 0.7075 | 0.7784 | 0.0710 | -0.3321 | 0.5543 | 0.5995 | 0.667 | 1.0000 | Uncertain | fraction |
| TL-O2-01 | switches_per_10ns | 4.0667 | 3.3333 | -0.7333 | -5.5333 | 4.0533 | 0.6056 | 0.667 | — | Uncertain | transitions per 10 ns |
| TL-O2-01 | lowF_area_nm2 | 1.1284 | 0.9543 | -0.1741 | -0.9682 | 0.5468 | 0.6406 | 0.667 | — | Uncertain | nm^2 |
| TL-O2-01 | effective_bins | 141.39 | 127.85 | -13.546 | -92.159 | 64.650 | 0.6488 | 0.556 | — | Uncertain | effective bins |
| TL-O2-01 | mode_rg_nm | 2.5553 | 2.4133 | -0.1420 | -0.3786 | 0.0946 | 0.8683 | 0.889 | — | Exploratory | nm |
| TL-O2-10 | compact_occupancy | 0.7075 | 0.8336 | 0.1262 | -0.2981 | 0.6289 | 0.6763 | 0.778 | 1.0000 | Uncertain | fraction |
| TL-O2-10 | switches_per_10ns | 4.0667 | 3.0667 | -1.0000 | -6.6000 | 4.7333 | 0.6331 | 0.556 | — | Uncertain | transitions per 10 ns |
| TL-O2-10 | lowF_area_nm2 | 1.1284 | 0.6756 | -0.4527 | -1.2238 | 0.1393 | 0.9060 | 0.889 | — | Supported | nm^2 |
| TL-O2-10 | effective_bins | 141.39 | 94.186 | -47.207 | -120.26 | 14.014 | 0.9122 | 0.889 | — | Supported | effective bins |
| TL-O2-10 | mode_rg_nm | 2.5553 | 2.3423 | -0.2130 | -0.5442 | 0.1183 | 0.8615 | 0.889 | — | Exploratory | nm |
| TL-versus-FL low-loading O2 similarity | overlap_2d | — | — | 0.4856 | 0.0295 | 0.6607 | 0.9813 | 1.000 | — | Strong | fraction |
| TL-versus-FL low-loading O2 similarity | jsd_bits | — | — | -0.5070 | -0.7365 | -0.0489 | 0.9827 | 1.000 | — | Strong | bits |

The table is restricted to compact occupancy, exchange, state-space breadth, modal Rg, and the low-loading O2 boundary-condition similarity contrast discussed in the main text.

**Figure S1.**
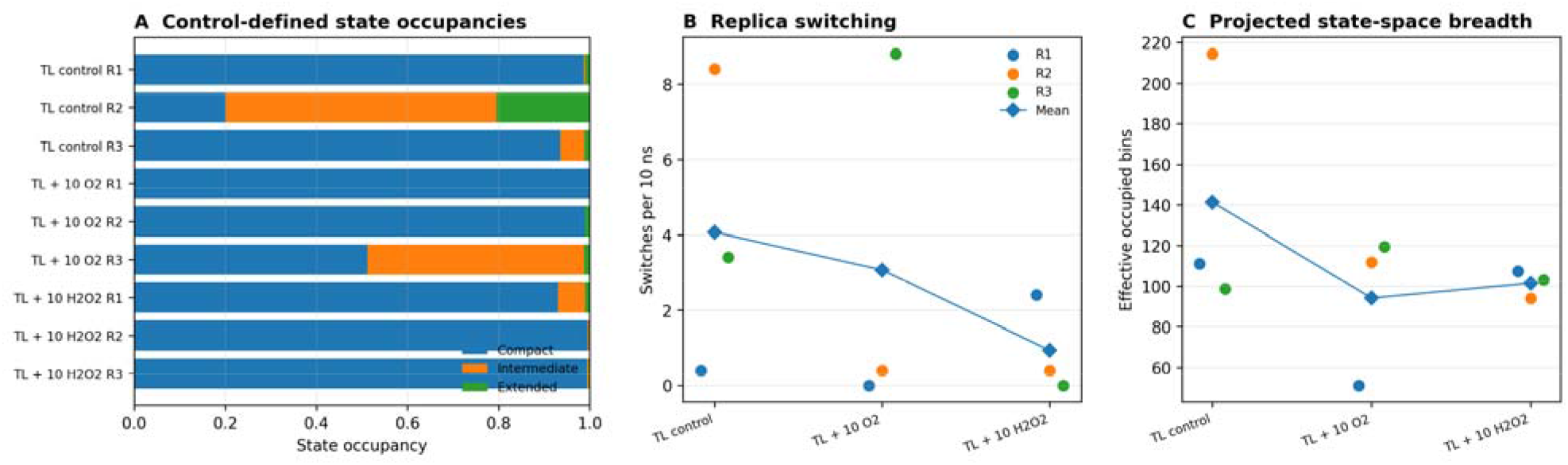
Replica-level heterogeneity of the high-loading tethered landscapes. **(A)** Control-defined compact, intermediate, and extended occupancies. **(B)** State switching per 10 ns. **(C)** Effective occupied-bin count as a projected state-space breadth descriptor. One TL + 10 O2 replica retained substantial intermediate-state occupancy and frequent exchange, whereas TL + 10 H2O2 was more consistently compact and low-exchange across replicas.

**Figure S2.**
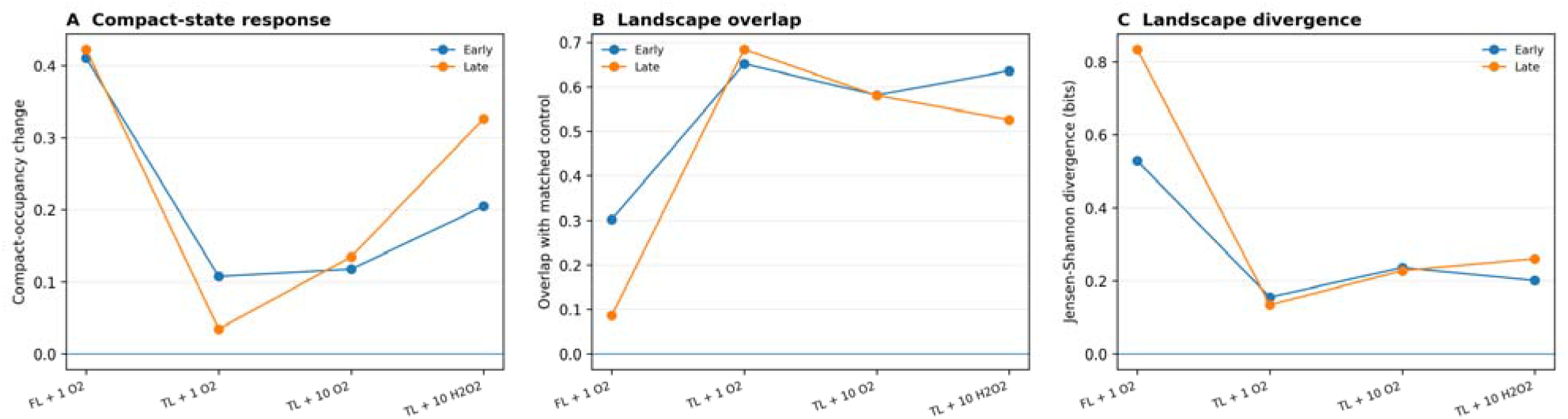
Early/late trajectory-window sensitivity of the principal landscape responses. Compact-occupancy changes, treatment-control overlap, and Jensen-Shannon divergence retained the main qualitative interpretation: FL + 1 O2 remained strongly displaced from control, TL + 1 O2 remained comparatively control-like, and high-loading TL O2 and H2O2 responses were directionally stable.

**Figure S3.**
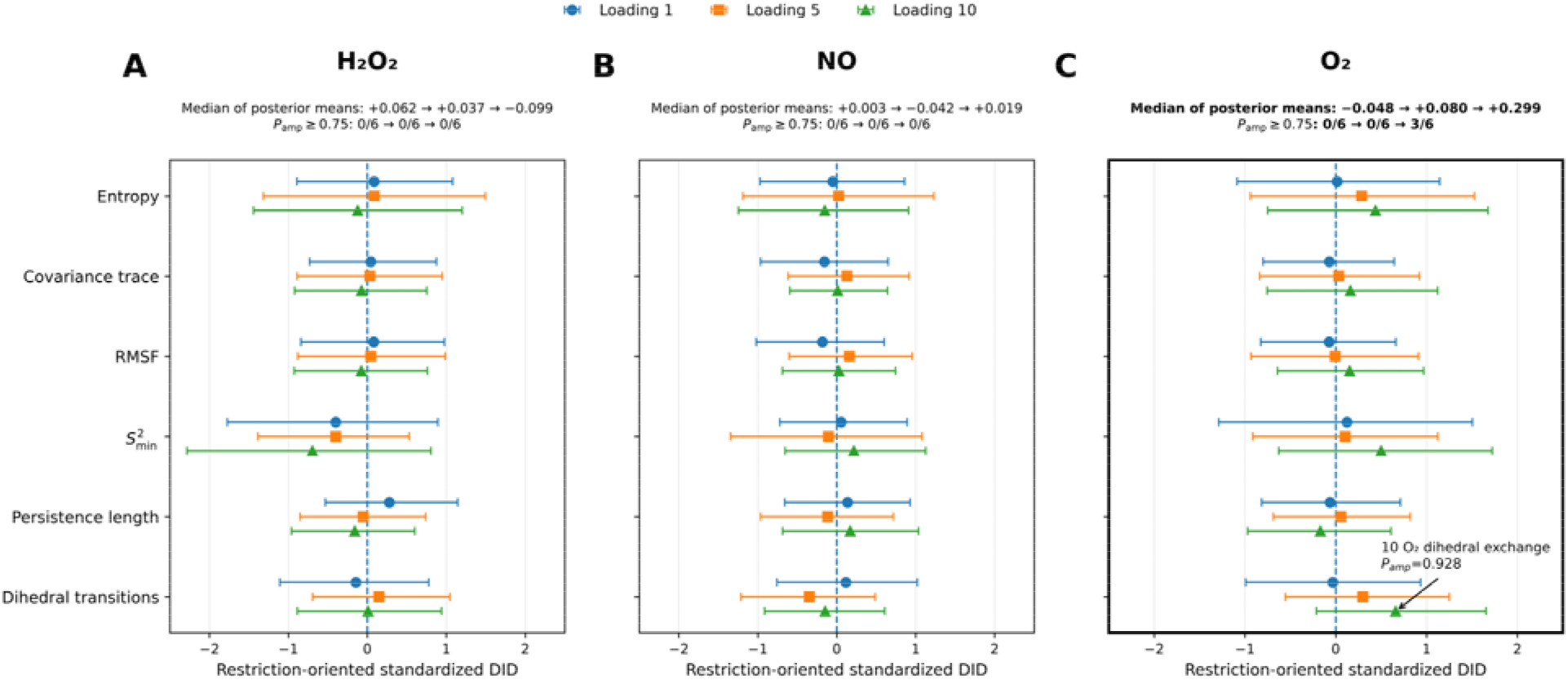
Posterior boundary-condition interactions show the strongest directional signal at high. **O**F **loading.** Points are posterior standardized DID means and horizontal lines are central 95% credible intervals. Positive values indicate amplification of restriction by tethering; negative values indicate attenuation or reversal. Circles, squares, and triangles denote loadings of 1, 5, and 10 molecules. Values above each panel summarize the median of the six posterior means and the number of metrics with Pamp ≥ 0. 75.

*The complete set of contiguous residue-level candidate regions underlying main Figure 4 is provided in Source_Data_File_3_Residue_Candidate_Regions data set in Source Data File*.

The source-data file reports all candidate regions, including opposite-direction responses, to avoid selective presentation. These are contiguous replica-robust regions and should not be interpreted as independently significant residues. Full residue-wise profiles remain part of the complete analysis archive.

*Source Data File. Condition-level inputs underlying the main Figure 5 association analyses are provided in Source_Data_File_4_Figure5_Condition_Data data set in Source Data File*.

The source-data file contains the integrated restriction index, its six component descriptors, intramolecular H-bond-count effects, and per-molecule energetic components for all 18 single-species treatment conditions. Energetic values retain the units reported in the manuscript (cal mol^-1 per species molecule).

**Table S5.** Condition-level mechanistic associations used in main Figure 5.

| Analysis | Statistic | Estimate | p value | 95% CI low | 95% CI high | Adjustment |
| --- | --- | --- | --- | --- | --- | --- |
| IRI vs change in intramolecular H-bond count | Spearman rho | 0.7152 | 0.0008 | 0.3347 | 0.9123 | None |
| IRI vs net attractive strength per molecule | Spearman rho | -0.4118 | 0.0895 | — | — | None |
| IRI vs net attractive strength per molecule | Partial Spearman rho | -0.2523 | — | -0.8215 | 0.4964 | log loading, species, mechanical state |

Condition-level associations use the 18 single-species treatments as independent observations. Bootstrap intervals were obtained by condition-level resampling.

Across the complete residue-level analysis, no global energy-dynamics association survived within-family multiple-comparison adjustment. Only five prespecified regional alignments met the energetic-colocalization criteria:

**Table S6.** Prespecified residue-level energetic alignments that met the colocalization criteria.

| Category | Contrast | Metric | Start_Residue | End_Residue | Dynamic_Direction | Dynamic_Classification | Dominant_Favorable_Component | Alignment |
| --- | --- | --- | --- | --- | --- | --- | --- | --- |
| Tethered single species | TL-H2O2-05 | RMSF | 654 | 657 | Restriction | Supported | electrostatic | Attractive hotspot overlaps restriction |
| Tethered single species | TL-H2O2-10 | S2Min | 668 | 670 | Restriction | Supported | electrostatic | Attractive hotspot overlaps restriction |
| Tethered single species | TL-NO-10 | RMSF | 635 | 637 | Restriction | Exploratory | vdw | Attractive hotspot overlaps restriction |
| Tethered single species | TL-O2-01 | S2Min | 637 | 639 | Restriction | Exploratory | vdw | Attractive hotspot overlaps restriction |
| Tethering DID | DID-O2-01 | S2Min | 637 | 639 | Tethering amplifies restriction | Exploratory | — | Energetic shift and dynamical DID agree |

The complete residue-level correlation table, including circular-shift p values and within-family adjusted q values, should be supplied separately with the full analysis archive; it is not reproduced in the formatted SI.

## Source Data File

The accompanying source-data file contains the following four data sets: 3

- Source_Data_File_1_State_Matched_Effects
- Source_Data_File_2_Model_Diagnostics
- Source_Data_File_3_Residue_Candidate_Regions
- Source_Data_File_4_Figure5_Condition_Data

## Notes

### Competing Interest Statement

The authors have declared no competing interest.

### Summary of Updates

This version of the manuscript has been revised to update the following. The title, abstract, introduction, and discussion have been rewritten to reflect an expanded study of noncovalent remodeling of the ABCB1 linker by selected plasma-associated molecular species. The simulation design now includes both free and structurally tethered linker models to distinguish responses dependent on unrestricted terminal motion from those retained under mechanical constraints. The expanded dataset comprises 60 independent 100 ns trajectories covering hydrogen peroxide, nitric oxide, and molecular oxygen at three molecular loadings, with matched controls. The statistical framework has been revised to include state-matched standardized effect sizes, Bayesian difference-in-differences analysis, replica deletion checks, and trajectory-window sensitivity analyses. New projected conformational landscapes characterize state populations and switching, while expanded residue-level analyses identify spatially heterogeneous responses. The results now distinguish the low-loading oxygen response that is largely suppressed by tethering from selected high-loading hydrogen peroxide and oxygen responses that remain detectable. Interaction-energy analyses have also been revised to show that total species-linker attraction does not reliably predict conformational remodeling, whereas intramolecular hydrogen-bond reorganization is more closely associated with conformational restriction. Methods, figures, figure legends, and references have been updated. The discussion of limitations has been expanded to clarify the reduced-model assumptions, sampling uncertainty, and restriction to noncovalent interactions. Conclusions have been revised to distinguish observed linker remodeling from untested effects on transporter activity, drug efflux, and reversal of multidrug resistance.

